# Development and validation of a respiratory syncytial virus multiplex immunoassay

**DOI:** 10.1101/2023.08.30.555534

**Authors:** Patrick Marsall, Madeleine Fandrich, Johanna Griesbaum, Manuela Harries, Berit Lange, RESPINOW study consortium, Stephanie Ascough, Pete Dayananda, Christopher Chiu, Jonathan Remppis, Tina Ganzenmueller, Hanna Renk, Monika Strengert, Nicole Schneiderhan-Marra, Alex Dulovic

## Abstract

Respiratory syncytial virus (RSV) is one of the leading causes of severe respiratory disease in infants and adults. RSV exists as two subtypes A and B, which co-circulate throughout the season, although one will usually become dominant. While vaccines and monoclonal therapeutic antibodies either are or will shortly become available, correlates of protection remain unclear. For this purpose, we developed an RSV multiplex immunoassay that analyses antibody titers towards the post-F, Nucleoprotein, and a diverse mix of G proteins. Technical and clinical validation showed outstanding performance, while methodological developments enabled identification of the subtype of previous infections through use of the diverse G proteins for approximately 50% of samples. As a proof of concept to show the suitability of the assay in serosurveillance studies, we then evaluated titer decay and age- dependent antibody responses within population cohorts. Overall, the developed assay shows robust performance, is scalable, provides additional information on infection subtype, and is therefore ideally suited to be used in future population cohort studies.

**Importance:** Although respiratory syncytial virus (RSV) is endemic and re-infections are common and harmless to the majority of the population, it is a leading cause of hospitalization in young children, the elderly, or immunocompromised individuals. A better characterization of RSV immunology and spreading dynamics is thus critical for preparedness, especially when interventions aiming to mitigate other diseases (e.g., COVID-19) disturb its endemic cycles. This requires high-throughput information-dense assays. We therefore developed a bead-based multiplex immunoassay that allows measurements of antibodies against multiple RSV antigens simultaneously. We identified antibodies which were strong indicators of previous infection, while others allowed identification of the subtype of the previous infection. The assay itself was shown to be robust and scalable, making it ideal for to keep track of the temporal variation RSV immunity profiles within the population.

## Introduction

Respiratory syncytial virus (RSV) is the leading cause globally of acute lower respiratory tract infections in infants [1–3], and is frequently the cause of pneumonia and subsequent hospitalization and mortality in older and immunocompromised adults [1, 4, 5]. As a negative sense, single stranded RNA virus, the RSV genome encodes for 11 proteins [6], of which the F and G glycoproteins induce the neutralizing antibody response [7, 8]. While the F protein is highly conserved among viral variants [9], the G protein shows high diversity with corresponding differences in monoclonal antibody reactions, resulting in two antigenic subtypes, A and B [10–12]. Recent genome sequencing revealed a wide variety of RSV genotypes, with a 2017 analysis identifying 11 RSV-A and 23 RSV-B genotypes [13]. While both A and B subtypes commonly co-circulate, one is usually predominant within a season [14]. However, it remains unclear whether one subtype causes more serious disease courses than the other, as studies identifying higher clinical severity have been published for both A and B subtypes [15–20]. Reinfections with RSV are common throughout life, with most individuals experiencing their first infection by the age of two [21], although this has likely been altered by the COVID-19 pandemic where population-wide non-pharmaceutical intervention (NPI) measures drastically altered the respiratory virus seasons [22]. For decades the only market approved product for paediatric immunoprophylaxis was the monoclonal antibody Palivizumab. Recently, however, not only has the monoclonal Nirsevimab been approved for pediatric use, but GSK’s and Pfizer’s subunit-based vaccines Arexvy [23] and Abrysvo [24] have both received Food and Drug Administration (FDA) approval for use in the elderly. Despite this, correlates of protection remain poorly defined.

Understanding RSV immunity and how it changes over time is critical to thereby assess potential future population dynamics, especially considering how these changed throughout the pandemic. This is only possible through assays that enable a deeper immune response profiling. Multiplex immunoassays in contrast to ELISAs offer the ability to measure antibodies towards an unlimited number of antigens simultaneously, making them a time-, sample- and cost-saving equivalent and suitable for use in epidemiological or vaccine studies. Therefore, we developed and validated an RSV multiplex immunoassay, which includes the post-F, Nucleoprotein, and diverse mix of G proteins as target antigens. As the assay is planned to be used to screen epidemiological cohorts, we orientated towards profiling G antibodies, as a way of identifying subtypes of previous infections.

## Results

### Improved G protein antibody detection through Anteo Coupling

All RSV antigens were coupled using either EDC/s-NHS or Anteo (see methods for details) in a variety of concentrations to determine optimal performance. While most antigens were unaffected or showed minimal changes in response to these different methods/concentrations in mean fluorescence intensity (MFI), there was a significant improvement in G protein performance when Anteo coupling was used (**Figure 1**). Compared to classical EDC-sNHS coupling, Anteo coupling resulted in significant increases in MFI values (all p<0.001), with subtype A G proteins increasing by 7.3 - 45.6x fold and subtype B G proteins increasing by 1.5 - 5.4x fold (**Figure 1**).

**Figure 1.**
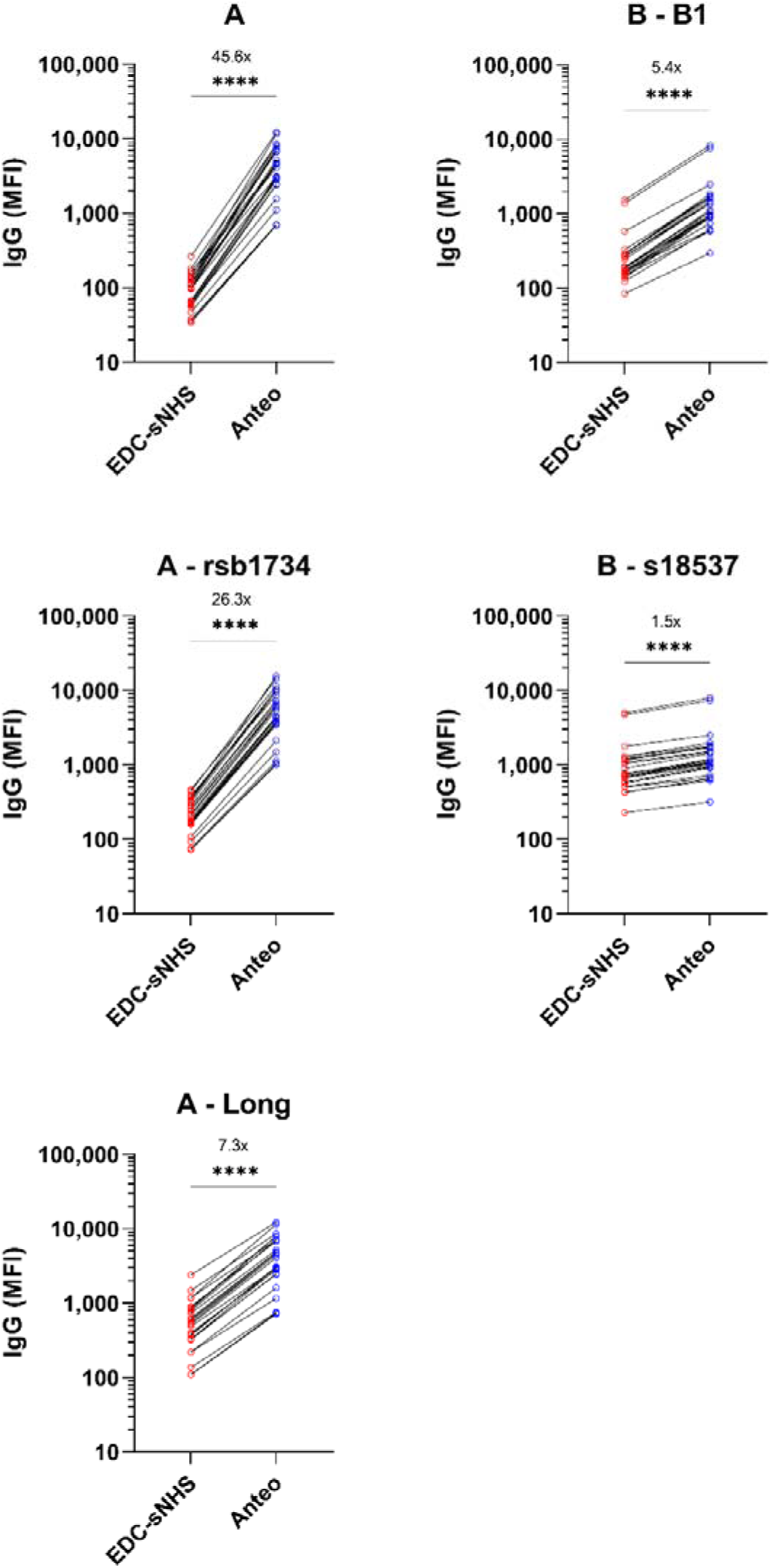
Improvement in G glycoprotein antibody binding with alternative coupling method. G glycoproteins were coupled using EDC-sNHS and Anteo (see methods for further details) at a range of different concentrations to determine the optimal conditions. Line graphs (a-e) showing change in antibody titer (IgG) for the different G proteins. The subtype and specific strains used are indicated in the title of each panel. Wilcoxon matched-pairs signed rank test was used to assess whether this change in titer was significant or not. **** indicates a p- value <0.0001. The mean change in antibody titer between EDC-sNHS and Anteo is included within each panel.

### Technical Assay Validation

Having identified the appropriate coupling conditions and concentration for each antigen, we assessed assay performance through technical validation. Five reference sera were serially diluted from 200 in four-fold steps to 3,276,800 to assess dilution linearity (**Figure 2a, Supplementary Figure 1**), identifying that our assay has a 3-log linear range corresponding to 250-25,000 MFI. All samples showed good parallelism for all antigens except the inactivated full virus. Inter- and intra-assay variability was exemplary, with low coefficients of variability (%CVs) for all antigens expect the full virus, which was therefore removed from the assay panel (**Figure 2b and c, Supplementary Table 1**). To evaluate antigen specificity and prozone effects, we spiked a post-F monoclonal antibody into both assay buffer and RSV- negative serum in a dilution series. Percentage recovery across all dilution factors met FDA bioanalytical guidelines (<15%) (**Figure 2d**). Antigens showed high specificity, with changes in antibody titer in response to spike in found only on the post-F beads at all dilution factors examined (**Figure 2e, Supplementary Figure 2**).

**Figure 2.**
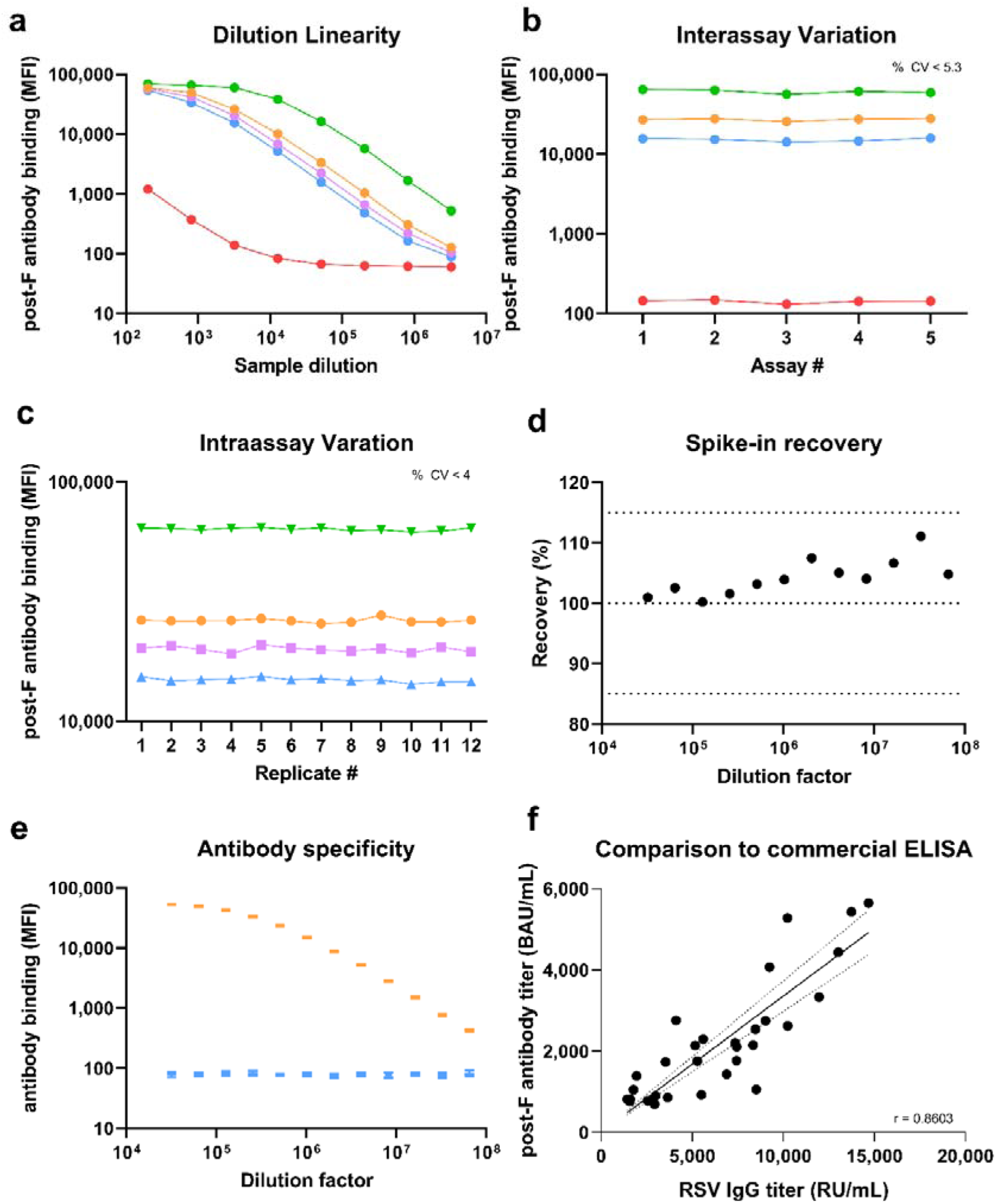
Technical validation of RSV multiplex immunoassay. Several parameters were assessed for technical validation of the RSV multiplex immunoassay. (a) Dilution linearity was assessed in five validation samples from DF200 – 3276800. Linearity within the assay corresponded to a range of 250-25000 MFI. Linearity for other antigens within the assay are included as Supplementary Figure 1. Inter- and Intra-assay variation (b and c) were assessed in four validation samples at DF3200. For Interassay variation, samples were measured in triplicate across five independent experiments, means of the triplicate for each sample is shown. %CV was <5.7% for all samples. For Intraassay variation, samples were measured in triplicate (technical replicates), 12x on a single plate. The mean of the technical triplicate for each of the 12 biological replicates is shown. %CV was <4%. CVs for Inter- and Intra-assay variation for all other antigens is included as Supplementary Table 1. To assess prozone effects (d), a monoclonal post-F antibody was spiked at various concentrations (DF32000-6.5x10^7^) into either 8 negative serum samples or assay buffer with percentage recovery calculated. Mean %recovery for the 8 samples is shown with 100% indicating no difference between serum and assay buffer. Standard bioanalytical margins for successful recovery (85% and 115%) are indicated on the panel. To confirm antigen specificity, binding titer for the post-F, N and G was assessed for the same antibody. Binding responses were found only on the post-F antigen bead (orange) and not on any others (G rsb1734 shown in blue as an example). Binding responses for the N is included as Supplementary Figure 2. Lastly, the RSV multiplex immunoassay performance was compared to a commercial RSV ELISA (see methods for details), with linear regression and Spearman r used to calculate agreement between the two assays. The correlation was highly significant (p<0.0001).

To confirm that the assay was not affected by its multiplex format, we evaluated assay performance as both monoplex and multiplex (**Supplementary Figure 3**). All antigens showed strong and significant correlations between monoplex and multiplex for all antigens (all Spearman’s correlation coefficients between 0.92 and 1.00, all p<0.001), confirming that the multiplex nature of the assay had no influence upon performance. Lastly, we compared our assay performance to a commercial anti-RSV IgG ELISA (**Figure 2f**), again identifying a strong and significant correlation between both assays (Spearman r = 0.86, p<0.001). This confirms that our assay performs at least as well as a comparable routine lab assay.

### Clinical Assay Validation

As RSV antibodies are widespread within the population, the assay was instead validated using samples from a challenge study, focusing on whether the assay could accurately detect changes in titer in response to infection. For samples from patients who went through an RSV infection, post-F and N antibodies significantly increased from day 0 to day 28 by 25.1 % (IQR=7.0-109.4%, p=0.02, **Figure 3a**) and 81.4% respectively (IQR 38.6-126.6%, p<0.001, **Figure 3c**). In contrast, control samples of uninfected individuals had non-significant changes of –0.8% (IQR –7.8 - +26.9, p=0.89, **Figure 3b**) and –0.1% respectively (IQR –7.7 - +18.2, p=0.74, **Figure 3d**). This increase in titer among infected individuals was also present at day 180, with an overall increase in titer from day 0 of 28.7% (IQR 2.1 - 55.5, p=0.03, **Figure 3a**) and 65.4% (32.7 - 102.0, p=0.002, **Figure 3c**) for the post-F and N, respectively. No significant change was observed between day 28 and day 180 for either the post-F (p=0.90) or N (p=0.98). Overall, we found that 30% (6 of 20) of the infected samples did not mount a detectable increase in titer in response to infection within our assay. To confirm that this was not a failing in assay performance, we assessed all samples from clinical validation with the commercial ELISA, which also identified the same samples having no change in antibody titer (data not shown).

**Figure 3.**
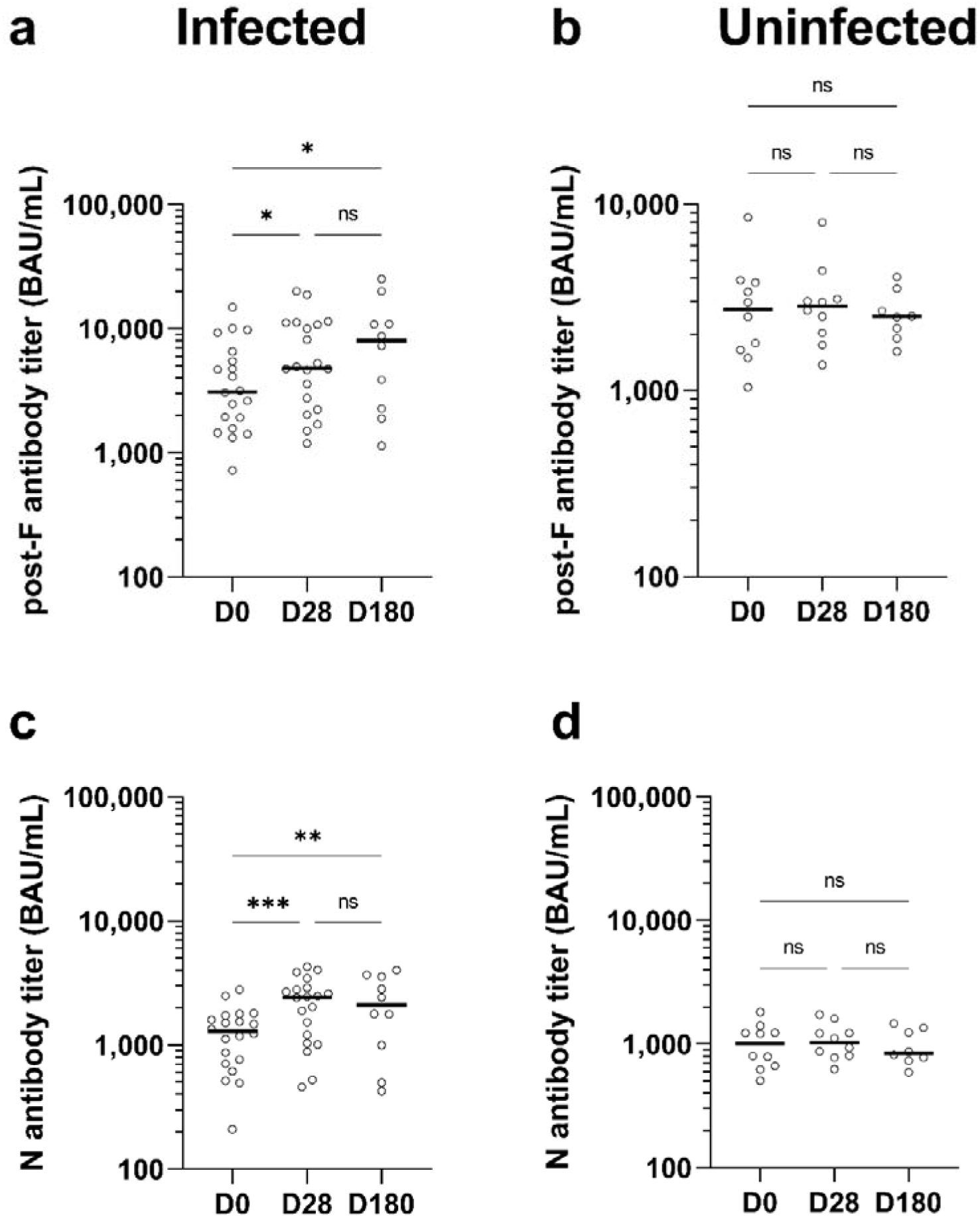
Clinical validation of RSV multiplex immunoassay. 20 samples from a human challenge study were used to evaluate the performance of the RSV multiplex immunoassay in detecting new infections. Samples were collected at day 0 prior to infection and at day 28 post infection (a and c). For some individuals, additional samples were collected at day 180. As a control, an additional group (n=10) who were not infected as part of the challenge study, had samples collected at the same timepoints (b and d). Changes in antibody titer were assessed for the post-F (a and b) and Nucleoprotein (c and d). Statistical differences in titer between timepoints was assessed using two-way ANOVA corrected with Tukey’s multiple comparisons test used for individual variances. ns indicates a non-significant p-value >0.05, * indicates a significant p-value <0.05, ** indicates a significant p-value <0.01 and *** indicates a significant p-value <0.001. Samples that were not showed no increase in titer in response to infection, were also analysed with the commercial ELISA which was in accordance.

### RSV antibody titer increases with age up to 5

Following technical and clinical validation, we assessed age-related RSV titers to gain an understanding into the overall landscape of RSV immunity in a cohort of 562 samples (derived from University Hospital Tübingen and MuSPAD, see Samples and Ethical Approval). Post-F and N titers increased with increasing age up to the age of 5 for post-F (median titer 886 BAU/mL) and N (median titer 0 BAU/mL) remaining stable at later ages (post-F median 2709 BAU/mL, N median 788 BAU/mL, **Figure 4a and b**). Titers themselves were highly individualistic, representing the pattern of continued re-infections with RSV. As expected, no negative samples were found after the age of 6, with the vast majority of negative samples coming from 1 or 2 year old individuals. To investigate this in more detail, we examined samples from a cohort of infants aged 12 to 36 months old at time of collection (University Hospital Tübingen). As all samples were collected during the first two years of the SARS-CoV-2 pandemic, exposure to RSV was less likely than in normal years. Overall, 32.8% (20 of 61) of samples were positive for both RSV post-F and N antibodies, whereas 55.7% (34 of 61) were negative for both post-F and N antibodies. The remaining 11.5% (7 of 61) were post-F positive only. To confirm that positive samples were a result of maternal antibodies, we analyzed IgA titers, identifying that 4 of 7 post-F positive only and 18 of 20 post-F and N positive samples had detectable IgA antibodies indicating a previous infection (**Supplementary Figure 4**). Interestingly, titers for individuals aged 65 or over, who are normally considered the most vulnerable group to RSV infection after young children, were slightly higher than young adults (>65 post F median titer 3099 BAU/mL, N median titer 1129 BAU/mL, 25-44 post F median titer 2477 BAU/mL, N median titer 726 BAU/mL, **Figure 4a and b**).

**Figure 4.**
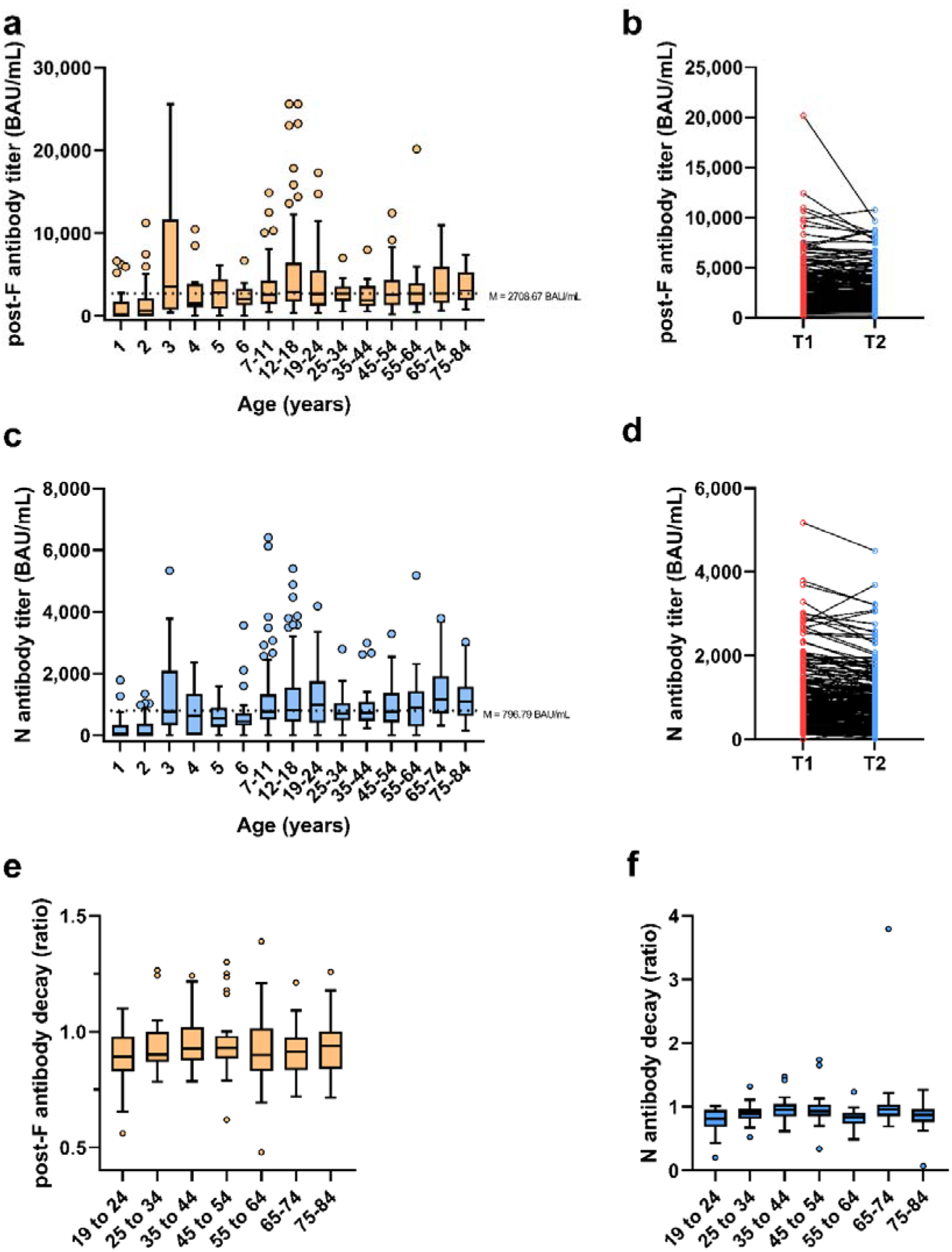
Age-specific pattern of RSV antibody titers and decay. RSV antibody titers towards the post-F and N were assessed using the RSV multiplex immunoassay in 562 individuals ranging from 1 to 84 years old. (a and c) box and whisker plots for post-F (a) and N (c) titer in across all ages, with boxes indicating the interquartile range and Tukey whiskers. Outliers are shown. Median titer from age 5 onwards is indicated. To assess antibody decay, line graphs showing longitudinal samples XX months apart from 172 individuals for the post-F (b) and N (d) antibodies. No samples from individuals considered to have been infected between sample collections were included (post-F and N titer increase both >25%). To evaluate whether changes in the rate of decay was linked to age, the same samples were stratified into age groups (e and f), with rate of decay for post-F (e) and N (f) calculated as change from T1 to T2.

### RSV antibody titer decay is limited

Having determined that RSV antibody titers appear stable from age 5 onwards, we next assessed long-term antibody presence by evaluating a longitudinal cohort of 190 individuals who donated samples in 2021 and 2022 (separated by 13-15 months) from the MuSPAD cohort, a German supraregional population-based cohort [25], adapted as an epidemic panel [26, 27]. To avoid including any individuals who had experienced infections between samplings, we excluded anyone who had a change in titer greater than 25% from 2021 to 2022 for both post-F and N (9% of samples). Overall, both post-F and N titers were highly stable, decreasing by 8.6% (0.5 – 15.4) and 9.5% (1.7 – 19.2),[26] respectively (**Figure 4c and d**). There was no significant effect of age upon rate of decay for either post-F or N (all p=0.99, **Figure 4e and f**).

### GA/GB antibody signatures allow identifying subtypes causing previous infection

While examining G antibody titers from the challenge study, we observed that half of the infected individuals had a greater increase in subtype A G antibodies than subtype B, and that no individual had a greater increase for subtype B (**Figure 5a**). To evaluate this further, we examined G antibodies within infants under 3 (**Figure 5b**), identifying that 85% (24 of 27) were heavily biased towards either subtype A or subtype B G antibodies. Among these, more than half (13 of 24) were positive for either subtype A or subtype B G antibodies only. Lastly, we evaluated GA/GB antibody signature within our longitudinal cohort to see how effective this signature was within real world samples. For samples that were classified as having been infected between sample collections (post-F and N titer increase by at least 25% each), 38.8% (7 of 18) of those had a GA/GB antibody signature that enabled classification as a previous subtype A or subtype B infection (**Figure 5c**).

**Figure 5.**
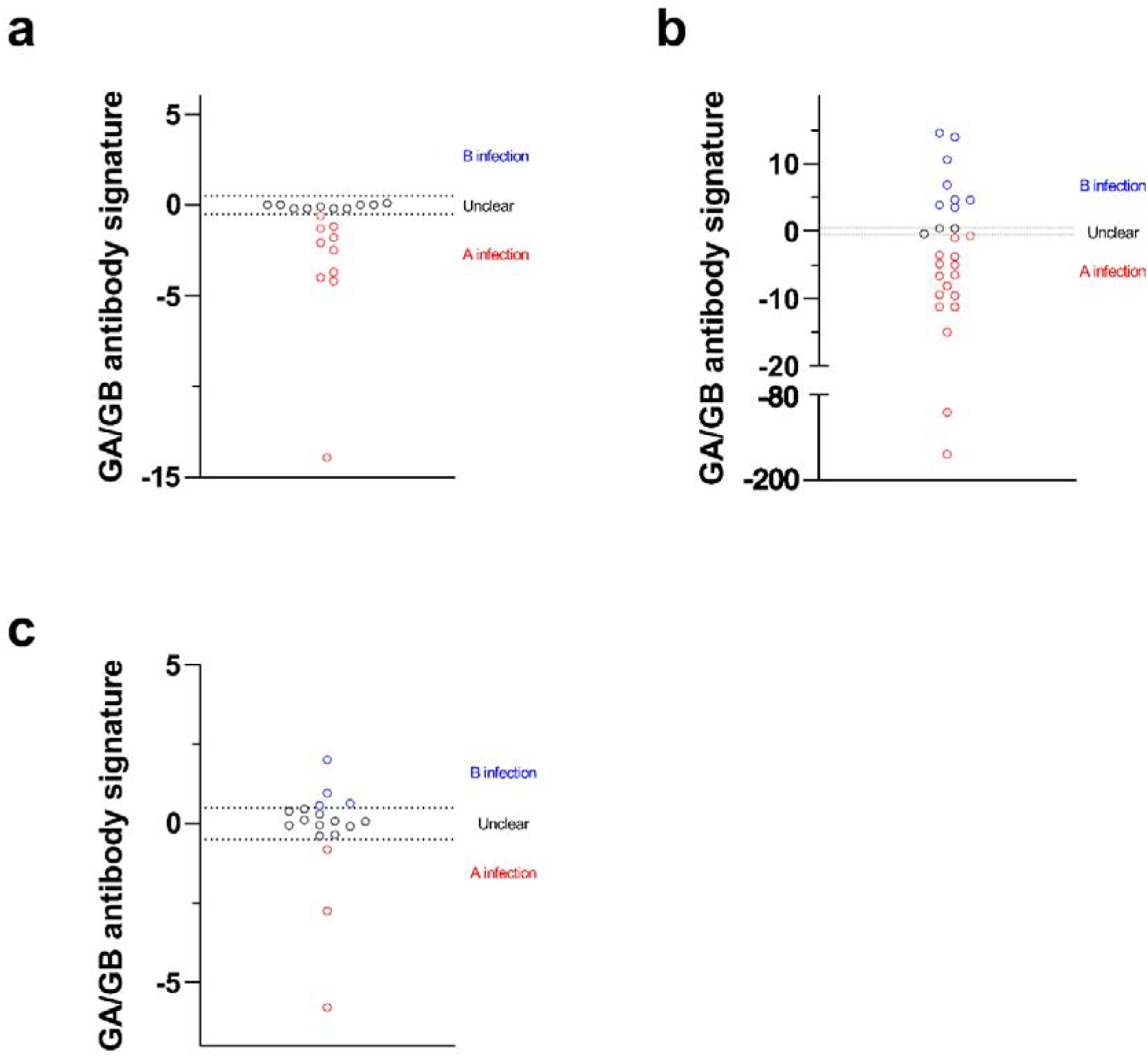
G antibody signature can be used to identify subtype of previous infection. GA/GB antibody signature was used where possible to evaluate the subtype of the most recent infection. A value greater than 0.5 was considered indicative of a previous subtype B infection, while a value greater than -0.5 was considered indicative of a previous subtype A infection. To validate this approach, samples from the challenge study who had been infected with Memphis-37 (subtype A) were evaluated (a), of which half of the samples showed a GA/GB antibody signature indicative of a previous subtype A infection, while the other half showed no specific signature. We then evaluated GA/GB antibody signature within children under 3 years old who were positive for RSV (b). Due to the limited number of previous infections, GA/GB antibody signatures were highly biased towards subtype A or subtype B, indicating the specific nature of the G antibody response. Lastly, we evaluated this within a population cohort (c) consisting of individuals aged 18 and over. For samples classified as having been infected between sample timepoints, a GA/GB antibody signature could be seen for 38.8% of samples.

## Discussion

While several RSV immunoassays have been previously described [28–30], this is the first to our knowledge that appears to actively use G antibody titers as part of the assay output. Critical to this was the increase in assay performance for the G antigens obtained through Anteo coupling, which enabled us to differentiate for some samples between historic subtype A and subtype B infections. A previous RSV multiplex immunoassay [30] that did include G antigens but only via EDC/s-NHS coupling, suspected that their low performance was either due to G proteins being less immunogenic or because their structure and low lysine content negatively affected conjugation. The improvements we saw from Anteo coupling suggest that G proteins are sufficiently immunogenic and that Anteo coupling may result in a more accessible orientation of the protein to the bead. Between subtypes, subtype A G antigen performance increased more than subtype B. Due to the low sequence homology between subtypes [12] and previous binding data suggesting shared epitopes [30], it may be that this difference is due to strain-specific epitope availability. As we previously have seen that Anteo coupling resulted in increased assay performance for other low performing antigens such as receptor binding domains (RBDs) from SARS-CoV-2 variants of concern [31–33], it will be interesting to assess other currently difficult to conjugate antigens and assess whether there are critical mutations/residues/structures for which Anteo coupling is optimal.

Overall, GA/GB antibody signatures could be determined for approximately a third of the samples we classified as having been infected between samplings in our longitudinal population cohort. While we obviously prefer to be able to differentiate for all samples classified as having been infected, this data still represents a major step forward and will be invaluable in epidemiological studies, allowing, e.g., direct comparisons between seasonal subtype prevalence within populations or regions of PCR-identified subtypes from severe hospitalized infections and assay-derived estimates of community transmission. Although we can validate this GA/GB antibody signature only for subtype A infections because of sample availability, our data from infants younger than 3 years suggest that we are correct in our approach, as shown by the single subtype G antibodies in some individuals. While it was not a focus of this study, it will be interesting to see whether strain-specific differentiation is possible, for which set of RSV-A strain Memphis-37 specific antigens are in development.

Our assay was highly sensitive and reproducible as demonstrated by the antigen specificity and exemplary CV values obtained. The three-log linear range enables the vast majority of samples to be measured at a single dilution factor. Since the international standard we used has identical arbitrary values for both RSV-A and RSV-B (1000 IU/mL) and no reference standard was available, we assigned 2000 IU/mL as a starting value for our assay standard. Based on the dynamic range of our assay, this meant that a standard curve of 0.0625 - 4 IU/mL could be recorded on each plate to enable conversion to BAUs. In the future, we will use an in-house developed reference comprising multiple serum samples covering a three-log range [34].

For clinical validation, instead of comparing to functional assays such as neutralization assays, we used samples from a human challenge study (see Samples and Ethical Approval) with a known infection status. As a result, we did not generate a cut-off value for positivity with our assay, although the individualistic nature of titers seen within our study cohort suggests this would be difficult to achieve. Critically, the assay was also able to detect negative samples among infants. Although the percentage of samples from infants, which were classified as negative by the assay was much higher than expected, it should be noted that these samples were collected during the pandemic at a time where RSV prevalence within communities was low [35, 36]. A recent monoclonal antibody study suggested that 25% of infants had undetectable neutralizing antibody levels at baseline [37].

In line with other publications [38], antibody titers for both post-F and N within our cohort increased until approximately age 5, after which they remained stable with increasing age. Interestingly for the two groups that are considered most at risk from RSV infections (under 5 and over 65), we saw opposing patterns for titers. As titer has been previously shown to correlate with neutralizing activity for RSV [30], further studies will be needed to elucidate why low titers in elderly individuals occur and appear to be less effective in at-risk groups than in younger counterparts. The overall decline we saw in RSV titer in a population-based cohort from Germany [25] is also similar to other reports assessing changes resulting from the pandemic [39]. While antibodies towards the F protein have been shown to correlate well with neutralizing activity [30], correlates of protection remain unclear [40], as does the impact of 2022-2023 RSV season where a surge in cases was reported.

Our assay offers several advantages over both classic RSV ELISAs and other serological assays. As a bead-based assay, it can be easily automated enabling high-throughput, while the multilog dynamic range means the majority of samples can be measured at a single dilution factor. Compared to single analyte assays like ELISAs, our assay not only offers a cost-, time- and material-saving alternative, but also provides additional information due to the antigen variety. The assay is also highly flexible and can be easily modified to measure other Ig isotypes (e.g., IgA, IgM) or IgG subtypes depending on sample material and study plan, while its modular format means that additional antigens (e.g., pre-F protein, more G proteins) can be added. Furthermore, our assay provides scalability for population-based studies. Maternal antibody transfer is a critical route for some of the RSV vaccines that are in development [41]. As RSV antibodies are placentally transferred during the third trimester and breast milk antibodies play a dominant role in neonatal mucosal immunity, it will be particularly interesting to assess the presence of RSV antibodies within breast milk.

The lack of cut-off within our assay suggests that it is more suited to use in longitudinal studies such as epidemiological screenings or time-derived vaccine or therapeutic studies. Unlike other RSV assays, we have not included the pre-F protein as an antigen as this was not available at the beginning of assay development. However, the flexible nature of our assay allows incorporating it once it becomes commercially available.

In conclusion, we have developed and validated an RSV multiplex immunoassay that shows strong stable performance, allowing for measurements of antibody titer towards a variety of RSV antigens, which is ideally suited for use in epidemiological or other longitudinal studies.

## Methods and Materials

### Samples and Ethical Approval

Samples from several different sources were used in this publication.

For technical assay validation, five reference sera were sourced from BEI Resources (#NR- 32832, contributed by NIAID and NIH), with an additional eight negative reference sera sourced from young children aged between 12 and 36 months who were born during the SARS-CoV-2 pandemic. To confirm their negative RSV status, we analyzed these samples with a commercial ELISA. The collection and use of these samples was approved by the Ethics Committee of Eberhard Karls University Tübingen and the University Hospital Tübingen under the ethical approval number 449/2022BO2 to Dr. Jonathan Remppis.

For clinical assay validation, 78 samples from 30 participants of an RSV human challenge study were used. Cohorts of healthy participants aged 65-75 years and 18-55 years were recruited in 2019-2023 and inoculated with 104 plaque-forming units (PFU) of RSV A Memphis-37 (M37). Serum samples were collected at day 0, 28 and 180 post-inoculation (p.i.). All participants had a matched day 0 and 28 p.i. sample, with 18 having a day 180 p.i. sample. Infection status was confirmed by N-gene specific qPCR, carried out on nasal lavage. Participants were regarded as RSV-infected following detection of virus on at least 2 consecutive days between day 2 and 10 p.i.. The study was approved by the Health Research Authority London-Fulham Research Ethics Committee (IRAS Project ID: 154109, REC References: 14/LO/1023, 10/H0711/94 and 11/LO/1826). All controlled human infection challenge (CHIM) studies were performed in accordance with ICH/GCP guidelines (US 21 CFR Part 50— Protection of Human Subjects, and Part 56—Institutional Review Boards). Prior to participation, written informed consent was obtained from all volunteers, who were free to withdraw at any time during the study.

To assess longitudinal antibody titers, 380 samples from 190 study participants from the MuSPAD [25] and RESPINOW [42] studies were used. All study participants donated two samples. Serum samples were originally collected as part of MuSPAD between 07/2020 and 08/2021, with a follow-up sample collected as part of RESPINOW between June and July 2022. To assess antibody titers in infants, 61 serum samples were collected from children aged between 12 and 36 months who were born just before (n = 35) and during the SARS-CoV-2 pandemic (n = 26). To assess the age-related structure of RSV antibody titers, we used 311 samples that were collected as part of a previous SARS-CoV-2 household exposure study [43], in addition to the samples stated above. Both, the original MuSPAD and RESPINOW sample, studies were approved by the Ethics Committee of Hannover Medical School (9086_BO_S_2020). The use of serum samples from young children in this study were approved by Ethics Committee of Eberhard Karls University Tübingen and the University Hospital Tübingen (449/2022BO2 and 293/2020BO2).

### Antigens and Antibodies

Antigens for assay development were purchased from Sino Biological and Aalto Bioreagents (see **Table 1** for full details). A commercially available F antibody (#11049-R302, Sino Biological) was also used during assay development.

**Table 1:**
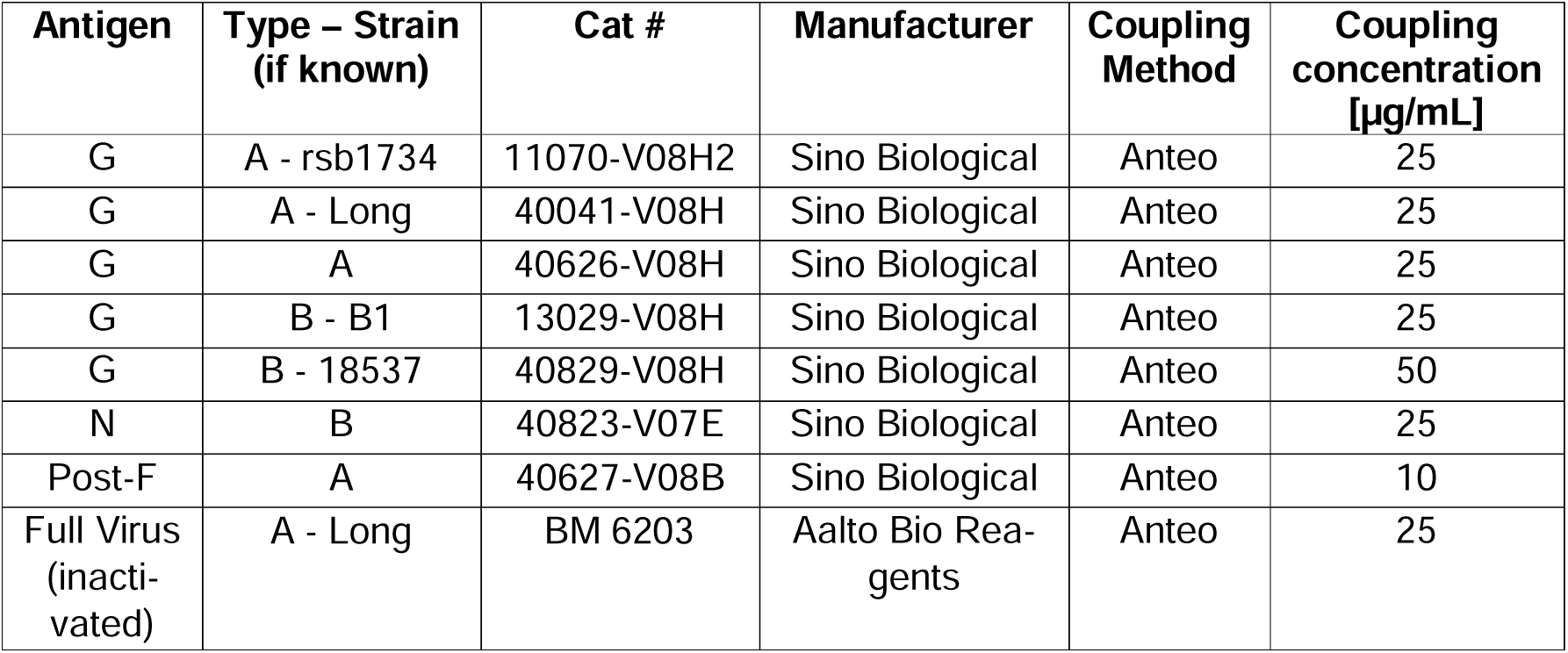
Antigens used during assay development including antigen, RSV subtype, strain (if known), the manufacturer and catalogue number as well as the final coupling method and concentration.

### Bead Coupling

All antigens were coupled to spectrally distinct populations of MagPlex beads (Luminex Corporation) using both EDC-sNHS and Anteo coupling at a variety of concentrations to determine the optimal coupling method and concentration for each antigen. EDC-sNHS coupling was performed as per the manufacturer’s instructions. Briefly, MagPlex beads were activated by using 1-ethyl-3-(3-dimethylaminopropyl) carbodiimide (EDC)/sulfo-N-hydroxysuccinimide (sNHS) chemistry. Bead stocks were incubated with 100 µL of an EDC/sNHS solution (100LJmM Na_2_HPO_4_, pH 6.2, 0.005% (v/v) Triton X-100 with a final concentration of 5 mg/mL for each reagent) for 20 min, washed twice with 250 µL of coupling buffer (500LJmM MES, pH 5.0, 0.005% (v/v) Triton X-100). Next, antigen solution in 500 µL of coupling buffer were added to the activated beads and incubated for 2 hours. Antigen-coupled MagPlex beads were washed twice with 500 µL of wash buffer (1x PBS, 0.005% (v/v) Triton X-100) and resuspended in 100 µL of storage buffer (1x PBS, 1% (w/v) BSA, 0.05% (v/v) ProClin). Bead stocks were stored at 4°C until required.

Anteo Coupling was performed using the AMG Activation Kit for Multiplex Microspheres (A- LMPAKMM-400, Anteo Technologies) as per the manufacturer’s instructions. Briefly, MagPlex beads were activated for 1 hour at room temperature in AnteoBind Activation Reagent, washed twice with conjugation buffer and then incubated with the antigen of interest for 1 hour at room temperature. Beads were then washed again twice with conjugation buffer and then blocked for 1 hour at room temperature with 0.1% (w/v) BSA solution. Following washing twice with storage buffer, the beads were resuspended in storage buffer and stored at 4°C until required.

### Commercial RSV ELISA measurements

60 samples were measured using EuroImmun RSV IgG ELISA (EI 2670-9601 G) as per the manufacturer’s instructions. Briefly, samples were diluted 1:100 in sample diluent buffer, added to individual wells of the plate and then incubated for 30 mins at 21°C, after which the plate was washed 3x with washing buffer. Enzyme-conjugate was added to each well and again incubated for 30 mins at 21°C. The plate was then washed again 3x with washing buffer, after which HRP-substrate was added to each well and incubated at 21°C. The reaction was stopped after 15 mins by the addition of stop solution. The plate was then measured using a BioTek ELX808 ELISA reader (Agilent) at 450nm and 620-650nm. Antibody titer (in RU/mL) was calculated using Gen5 software (version 1.10.8).

### RSV Multiplex Immunoassay

Individual bead populations were combined to generate a bead mix with a concentration of 500 beads/per ID/per well. Serum samples were thawed, diluted across two steps to 1:1600 in assay buffer [44] and then transferred into individual wells of a 96 half-well plate (Corning, Cat #3642). An equal volume of bead mix was then added to each well (final dilution factor 1:3200) and incubated on a thermomixer at 21°C for 2 hours at 750 rpm. Following this, the plate was washed 3x with Wash Buffer (1x PBS and 0.05% (v/v) Tween-20) using a magnetic plate washer (Biotek 405TS, Biotek Instruments GmbH). To detect bound IgG RSV antibodies, 3 µg/mL R-phycoerythrin-labeled goat anti-human IgG antibody (109-116-098, Dianova) diluted in assay buffer was added to each well and incubated for 45 mins, 750 rpm, 21°C on a Thermomixer. The plate was then washed again 3x to remove unbound antibodies, after which the beads were resuspended in 100µL wash buffer, shaken for 3 mins on a Thermomixer (21°C, 750rpm) and measured using an INTELLIFLEX-DRSE (Luminex Corporation) under the following settings: volume 80µL, count 50, gating 7000-17000. As a control and to enable conversion from MFI values to binding antibody units/mL (BAU/mL), the 1^st^ International Standard for Antiserum to RSV (16/284) was included a dilution series from 8 to 0.125 IU/mL. BAU/mL for each sample were calculated according to sample dilution factor using 7-parameter nonlinear regression as well as interpolation of the standard curve. If a sample did not fit within the linear range of the curve, it was measured at a higher dilution factor (upto 1:12800).

### Technical assay validation

Technical validation was performed according to the FDA guidelines for the validation of bioanalytical methods [45]. Technical measures of assay performance assessed were dilution linearity, interassay variance, intraassay variance, effect of multiplex format and determination of antigen specificity and cross-reactivity.

Dilutional linearity of the assay was determined using five different reference sera (all sourced from BEI resources) being two-fold serially diluted from 1:200 to 1:3276800 (**Figure 2a, Supplementary Figure 1**). Interassay variance was assessed by measuring four reference sera in triplicate across five assay plates. Experiments were performed across multiple days by multiple experimenters. Percentage of coefficients of variability (%CV) as measure for variance for each individual antigen was then determined (**Figure 2b, Supplementary Table 1**). For intraassay variance, four reference samples were measured in 12 biological replicates on a single plate. This was then repeated three times, with %CV then calculated for per antigen and per replicate (**Figure 2c, Supplementary Table 1**). To assess prozone effects from serum, a monoclonal post-F antibody was spiked in a dilution series (32,000 – 65,536,000) into 8 negative sera and assay buffer. Differences in MFI between the two-sample matrix were calculated as percentage recovery, with 100% indicating no difference between the two matrices (**Figure 2d**). To evaluate antigen specificity, antibody responses towards the sample monoclonal antibody were assessed for the post-F, G (rsb1734) and N (**Figure 2e, Supplementary Figure 2**). To identify cross-reactivity within the bead mix, 16 samples were measured using individual bead populations (monoplex) and the multiplex bead mix (**Supplementary Figure 3**), with differences in MFI between the monoplex and multiplex evaluated. Lastly, to compare assay performance to a commercial assay, a validation set of 30 samples was measured with multiplex RSV immunoassay and the anti-RSV IgG ELISA kit (Euroimmun). Results were correlated for the post-F glycoprotein with each other to verify specificity of both assays (**Figure 2f**).

### Clinical assay validation

Samples from the challenge study were measured with the RSV multiplex immunoassay and commercial RSV ELISA as described above.

### Data analysis

Binding Antibody Units per mL (BAU/mL) was calculated according to sample dilution factor using 7-parameter nonlinear regression as well as interpolation of the standard curve. Metadata was merged with analytical data in Excel, after which data analysis and visualization were performed in GraphPad Prism 9 (version 9.4.1). G antibody signature, as an indicator of subtype of the most recent infection, was analyzed by calculating the change in titer as a ratio for each G antibody titer, with mean changes in titer for subtype A and B then generated. The average of subtype A was subsequently subtracted from the average of subtype B, resulting in the GA/GB antibody signature. A GA/GB antibody signature >0.5 was considered indicative of an RSV-B infection, with a value <-0.5 indicating an RSV-A infection. The exact statistical test used is stated in each figure legend. Assessment of significant differences between groups was analyzed using a two-way ANOVA with Tukey’s multiple comparisons test. Linear regressions were used to evaluate correlations between ELISA- and RSV multiplex immunoassay-tested samples and between multiplex and monoplex assay performance, with Spearman r statistic used to estimate a rank-based association of two variables.

## Acknowledgements

We thank all participants supporting and endorsing the development of this multiplex assay. We are grateful to our colleagues in Plauen and Oldenburg for their assistance with aliquot preparation. We acknowledge the support of the Medical Research Council (G0902266), the Wellcome Trust (087805/Z/08/Z), the Kwok Foundation, and Medical Research Council (MRC) EMINENT Network (MR/R502121/1) which is co-funded by GSK. Infrastructure support was provided by the NIHR Imperial Biomedical Research Centre and the NIHR Imperial Clinical Research Facility. The views expressed are those of the authors and not necessarily those of the NHS, the NIHR or the Department of Health and Social Care.

