## Supplementary files for "Development and validation of a respiratory syncytial virus multiplex immunoassay"

**Supplementary Table 1: Inter and Intra assay variation for all other antigens in the RSV multiplex Immunoassay**

|  |  | Post-F - A | G - A | G – A, rsb1734 | G – A, long | Full virus – A, long | N - B | G – B, B1 | G – B, s18537 |
| --- | --- | --- | --- | --- | --- | --- | --- | --- | --- |
| Intraassay | Mean | 3.6 | 3.9 | 3.5 | 3.6 | 11.9 | 4.4 | 4.5 | 3.9 |
|  | Maximum | 4.0 | 4.7 | 4.1 | 4.3 | 15.3 | 5.7 | 5.0 | 4.3 |
| Interassay | Mean | 4.9 | 7.2 | 8.4 | 9.1 | 17.4 | 7.2 | 8.6 | 6.4 |
|  | Maximum | 5.3 | 9.4 | 11.4 | 11.5 | 23.1 | 9.2 | 14.0 | 7.4 |

%CVs for Intra- and Interassay variation for all antigens within the RSV multiplex immunoassay. For each antigen, the subtype and specific strain when known is indicated. Both the mean of the five reference samples and the single highest value (maximum) are shown.

**Supplementary Figure 1 – Dilution Linearity for other antigens in RSV multiplex immunoassay**

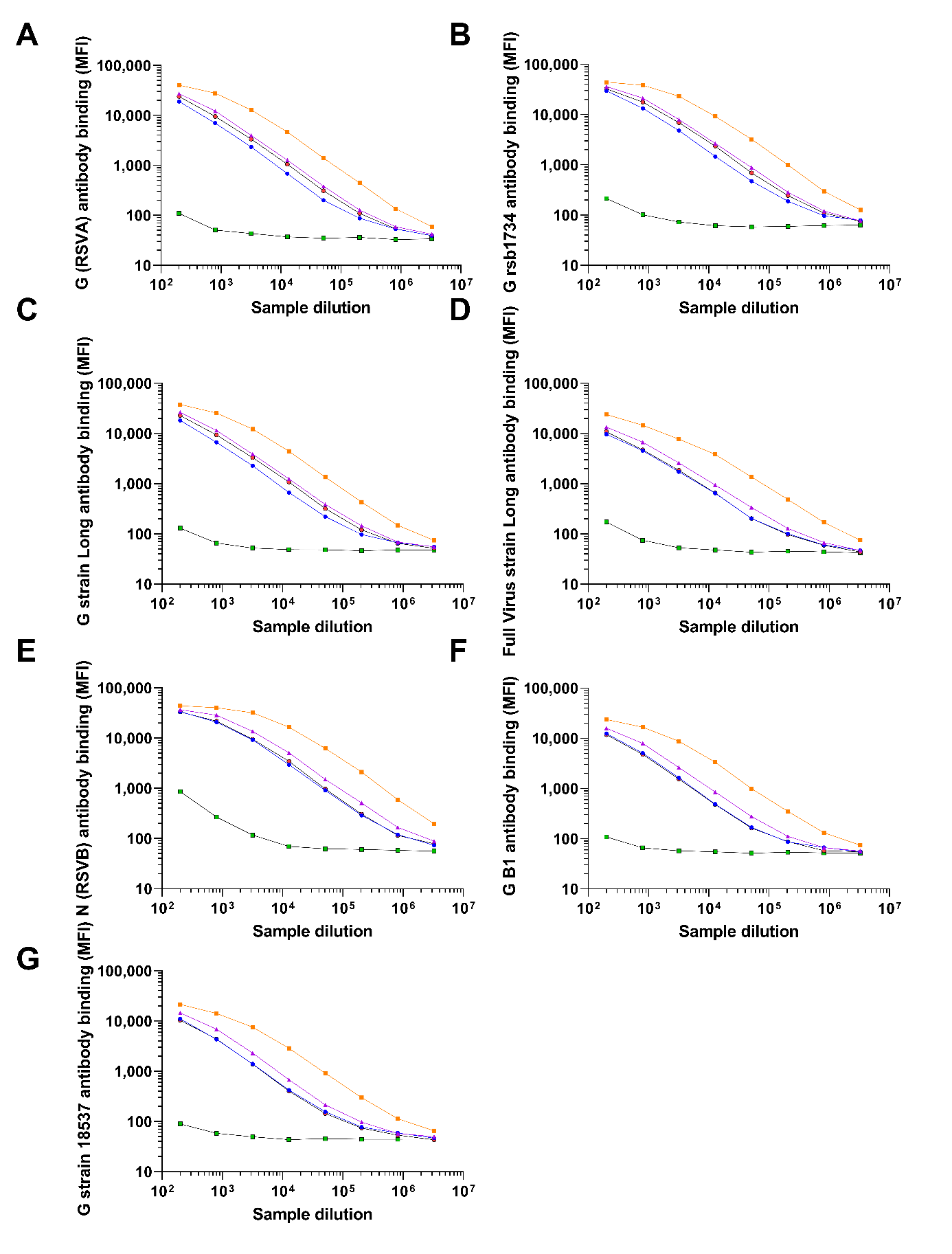

Dilution linearity for all antigens in the RSV multiplex immunoassay was assessed as part of technical validation. For this, five validation samples were measured from DF200 – 3276800, demonstrating a 3 log linear range from 250 to 25000 MFI.

**Supplementary Figure 2 – No change in N antibody binding following F-antibody spike in**

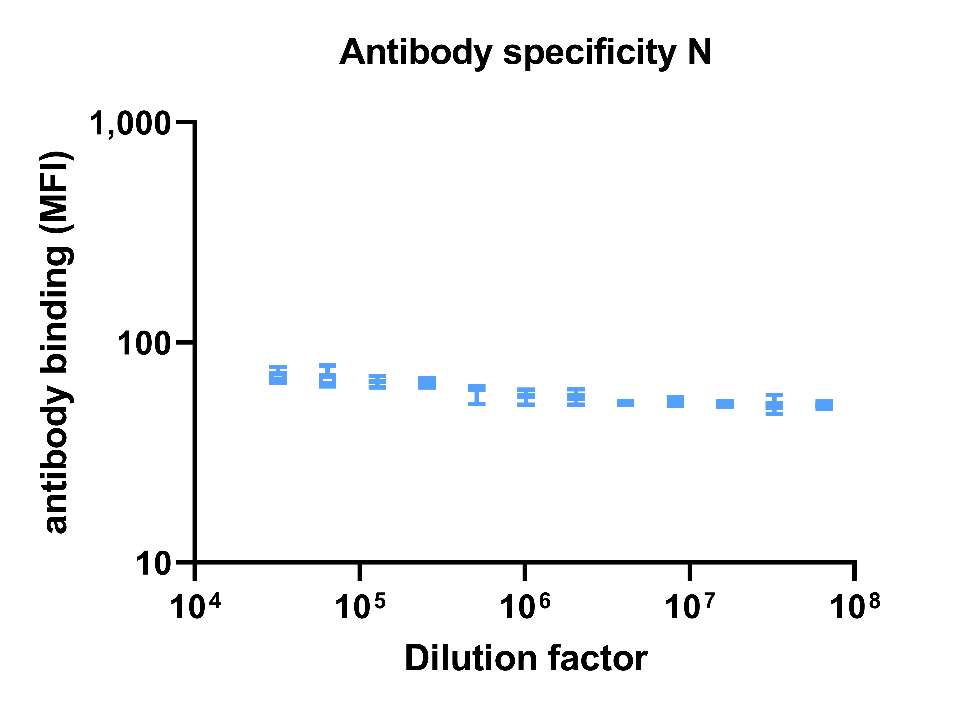

To assess antigen specificity, a monoclonal post-F antibody was spiked at various concentrations (DF 32000-6.5x10^7^) into assay buffer, with antibody binding towards the post-F, N and G analysed. Post-F and G are included in the main manuscript file as **Figure 2e**. There was no change in N titer in response to the antibody spike-in confirming antigen specificity.

**Supplementary Figure 3 – No change in assay performance as monoplex versus multiplex**

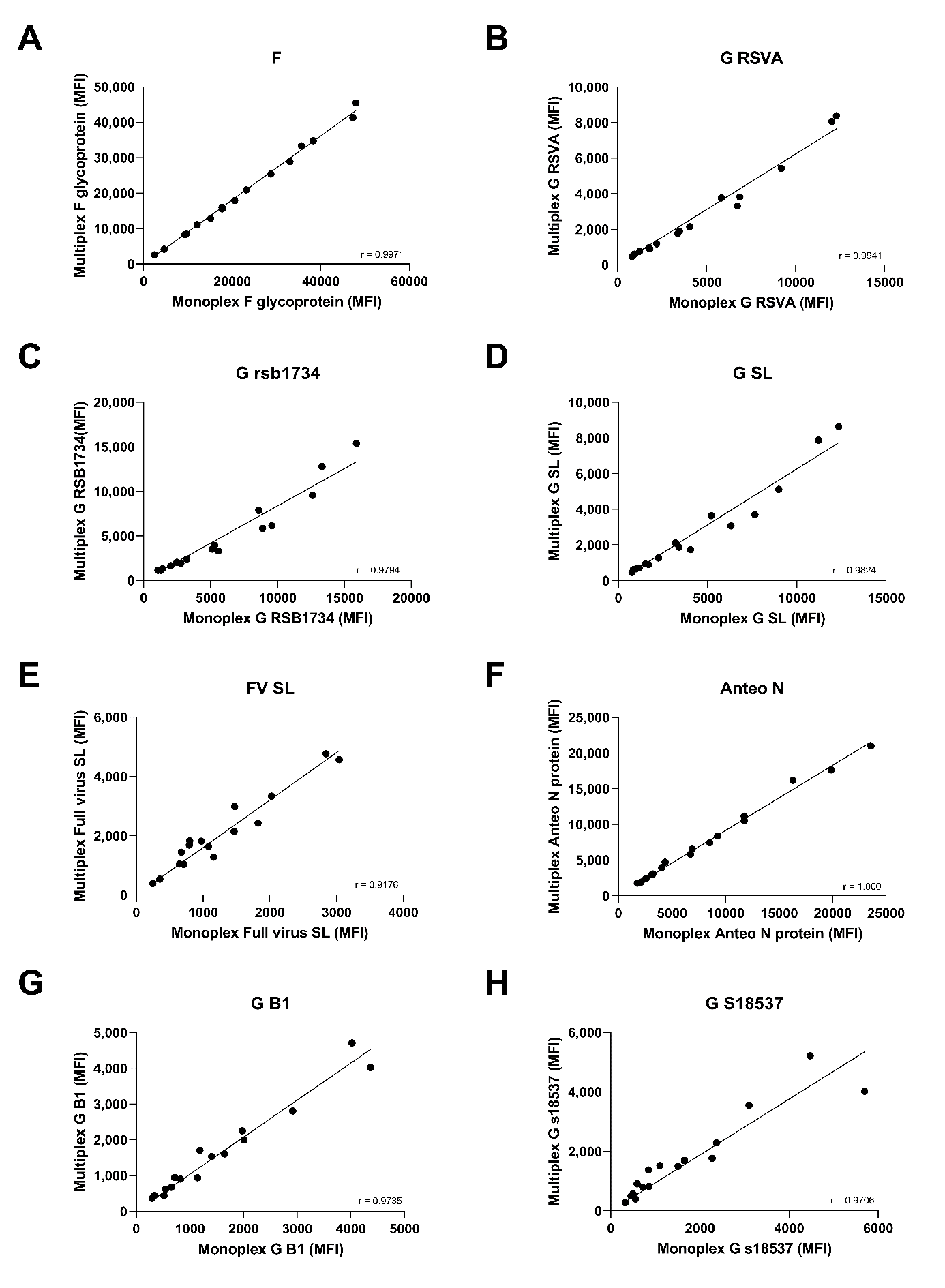

To confirm that antibody binding was not affected by the multiplex format, we assessed antibody binding in 16 samples for each antigen in both multiplex and monoplex format. Differences between the two assay formats were analysed using linear regression with Spearman correlation used to determine agreement. Spearman values for each correlation are included within each panel.

**Supplementary Figure 4 – IgA antibody presence in infant samples to confirm infection**

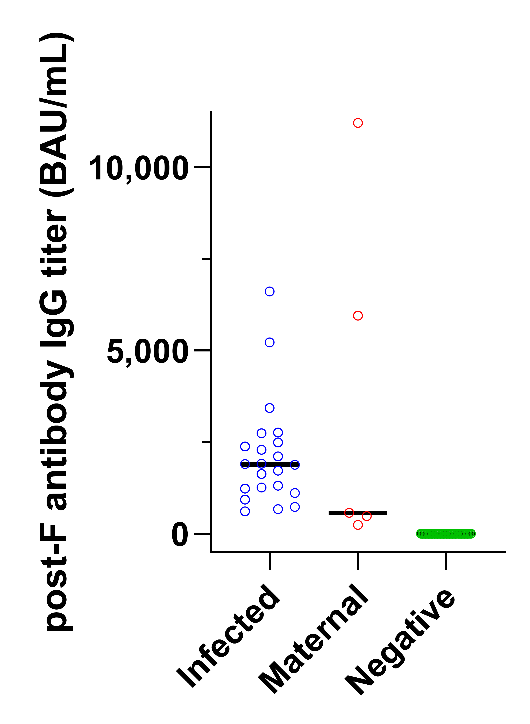

To confirm whether infected samples in infants had been truly infected or had antibody titers as a result of maternal transfer, we analysed post-F IgA titers. Samples which had been IgG negative were also IgA negative. For samples which were IgG positive, 5 of 27 had no detectable IgA indicating titers were as a result of maternal antibody transfer.

**Members and Contributors of RESPINOW study consortium**

Alex Dulovic, André Karch, Berit Lange, Carolina Klett-Tammen, Claudia Denkinger, Cornelia Gottschick, Daniel Junker, Daniel Wolffram, Felix Guenther, Isti Rodiah, Johannes Bracher, Laura-Inés Boehler, Lisa Koeppel, Manuela Harries, Melanie Schienle, Nicole Schneiderhan-Marra, Olga Hovardovska, Patrick Marsall, Philipp Dönges, Rafael Mikolajczyk, Sebastian Contreras, Torben Heinsohn, Ulrich Reinacher, Veronika K. Jaeger, Viola Priesemann
